# Ultrastructure of the nebenkern during spermatogenesis in the praying mantid *Hierodula membranacea*

**DOI:** 10.1101/2023.04.16.537100

**Authors:** Maria Köckert, Chukwuebuka William Okafornta, Charlice Hill, Anne Ryndyk, Cynthia Striese, Thomas Müller-Reichert, Leocadia Paliulis, Gunar Fabig

**Affiliations:** Experimental Center, Faculty of Medicine Carl Gustav Carus, Technische Universität Dresden, 01307 Dresden, Germany; Biology Department, Bucknell University, Lewisburg, PA 17837, USA

## Abstract

Spermatogenesis leads to the formation of functional sperm cells. Here we have applied high-pressure freezing in combination with transmission electron microscopy (TEM) to study the ultrastructure of sperm development in subadult males of the praying mantid *Hierodula membranacea*. We show the ultrastructure of different stages of sperm development in this species. In addition, we have applied serial-section electron tomography of the nebenkern to demonstrate in three dimensions (3D) that this organelle is composed of two interwoven segments that are connected by a zipper-like structure at opposing positions. Our approach will enable further ultrastructural analyses of the nebenkern also in other organisms.

## Introduction

The praying mantids (order Mantodea) have been used as a system for studying spermatocyte meiosis for over a century, with the first publication describing aspects of mantid male meiosis in 1897 (1). Unique features of chromosome behavior during both meiosis I and meiosis II have contributed greatly to our understanding of chromosome condensation, the balance of forces in spindles, and the spindle checkpoint(2–9). While many insights into spermatocyte meiosis have resulted from studies using praying mantids, there are few reports of other stages of spermatogenesis in mantids. The only survey of spermatogenesis in a praying mantid was performed by Williams(10), who studied this process in the mantid *Choeradodis rhombicollis*, showing drawings of different stages of spermatogenesis in fixed and stained specimens observed through a light microscope. More recently, electron microscopy of mantid sperm(11,12), and male reproductive anatomy(13) have also been studied, but a thorough survey of spermatogenesis including ultrastructural features in a praying mantis has not been published so far. Such a detailed study on the structure of cells in the testes of a mantid by both light and transmission electron microscopy (TEM) has the potential to reveal a great deal about sperm development in general, providing an additional insect order for comparative work. Because the cells are large and can lie flat on a slide for light microscopy, and have large, clearly visible organelles under any type of microscopy, they allow for detection of more difficult to see cytological details, thus informing previous studies in other organisms.

Some of the earliest studies of spermatogenesis in insects revealed the existence of a relatively large, layered structure made up of mitochondria, which was called the nebenkern(14,15). The nebenkerns of several insect species (Hemiptera, Heteroptera) were characterized by light microscopy(15,16) and later by electron microscopy(17,18), showing that the nebenkern forms at the end of meiosis II through the fusion of mitochondria. The nebenkern is composed of two halves separated by a furrow, with interdigitating layered membranes and thin bridges connecting the two halves(17). Further studies of spermatogenesis in *Drosophila melanogaster* revealed that this insect also forms a nebenkern during spermatogenesis that also displays a layered so-called “onion” structure, composed of two halves divided by a furrow(19,20). Examination of spermatogenesis in *D. melanogaster* allowed the merging of both light microscopical and ultrastructural data, and molecular genetics, showing that the initial fusion of the mitochondria to form the nebenkern requires intact microtubules, the dynamin-related protein Drp1, and components of the pink1/Parkin pathway(21). Formation of a functional and visibly wild-type nebenkern also requires other proteins that mediate mitochondrial fusion, like Fzo (an orthologue of the human mitofusins, Mfn1 and Mfn2), and the testis-specific ATP synthase subunit d paralog knon(22–24).

Previous studies in insects have revealed interesting membrane features associated with spermatogenesis, variations in flagellar structure, and variations in the structure and fusion of mitochondria associated with the formation of the nebenkern(12,17). The aim of this study is to show new information about spermatogenesis through an initial characterization of cells in the testes of the praying mantid *Hierodula membranacea*. Using state-of-the-art sample preparation techniques in combination with three-dimensional (3D) imaging such as high-pressure freezing and serial-section electron tomography, we show a survey of the steps of spermatogenesis in this species. Our study reveals interesting features of cellular structures during spermatogenesis, including the structure of the chromosomes in prophase I and the distribution of organelles in spermatids. In particular, we show the existence of the nebenkern in spermatids, revealing that the nebenkern of this species has two zipper-like membrane structures at opposing positions.

## Materials and methods

### Handling of subadult mantids

Subadult males of *Hierodula membranacea* (Giant Asian Mantis) were obtained from InsectSales.com (Port Angeles, Washington, USA) and Mantids & More (Mühlheim am Main, Germany). The authors verified the vendor’s identification according to previous publications(25,26). Mantids were fed house flies and crickets according to instructions from the vendors. To remove testes from subadult males, slits were made on both sides of the posterior of the abdomen, just anterior to the last abdominal segment. Testes were gently squeezed out of the slits, removed with tweezers and then placed in either Voltalef oil (Atofina) for phase contrast light microscopy or phosphate buffer (0.1 M, pH 7.4) for preparation for electron microscopy.

### Light microscopy

Live-cell imaging of subadult male testes was performed as described in Lin et al. (27). Briefly, spermatids at different stages of development were imaged across multiple focal planes using a Nikon Eclipse TS100 microscope (Nikon Instruments Inc., New York, USA) equipped with a 100x, 1.25 NA phase-contrast, oil immersion objective. Images were recorded using a View4K HD camera (Microscope Central) within the InFocus software (Microscope Central).

### Electron microscopy

#### High-pressure freezing, freeze substitution and resin embedding

For ultrastructural preservation of the male gonad of *H. membranacea*, high-pressure freezing in combination with freeze substitution was applied. Cryo-immobilization of the gonads was achieved by cooling the sample down to liquid nitrogen temperature (−196 °C), while exposing a pressure of approx. 2000 bar(28). In preparation for freezing, sample holders (type-A aluminum planchettes, Wohlwend, Switzerland) were pre-wetted with hexadecene (Sigma). For each freezing run, the 200 µm indentation of a planchette was then filled with 0.1 M phosphate buffer containing 10 % (w/v) polyvinylpyrrolidone (PVP, Sigma, MW 10,000(28–30)), and pieces of the dissected mantid testes were transferred to the cavity. The type-A planchette was then closed with the flat side of hexadecene-pre-wetted type-B planchette (Wohlwend, Switzerland). These ‘sandwiches’ were then immediately frozen using a high-pressure freezer (HPF Compact 03, Wohlwend, Switzerland) and stored in liquid nitrogen.

For subsequent freeze substitution, the cryo-immobilized samples were transferred under liquid nitrogen to a frozen ‘cocktail’, consisting of 0.1 % (w/v) uranyl acetate (Polysciences, USA) and 1 % (w/v) osmium tetroxide (EMS, USA) in anhydrous acetone (EMS, USA). The freeze substitution was performed with an automated freeze-substitution machine (AFS2, Leica, Austria). The samples stayed at -90 °C for one hour. The temperature was then raised to -30 °C with incremental steps of 5 °C per hour. The samples were kept at -30 °C for 5 hours. Afterwards, the temperature was raised again by 5°C per hour to 0°C, and the samples were kept for three additional hours until they were further processed.

#### Thin-layer embedding, serial sectioning and post-staining

After freeze substitution, samples were washed three times in pure acetone and infiltrated with resin as previously published(31). In brief, samples were gradually infiltrated with Epon/Araldite resin (one part resin : three parts acetone) for 1 h; 1 : 1 for 2 h; 3 : 1 for 2 h, and 100 % resin for 1 h, then 100 % resin overnight, then 100 % resin for 1 h and thin-layer(31). Resin-infiltrated samples were polymerized for three days at 60°C.

Selected samples were re-mounted on dummy blocks for ultramicrotomy(31). Serial thin (70 nm) sections for routine transmission electron microscopy and semi-thick (300 nm) sections for electron tomography were cut using an ultramicrotome (EM UC6, Leica Microsystems, Austria) equipped with a diamond knife (Diatome, Switzerland). Sections were collected on Formvar-coated copper slot grids and post-stained with 2 % (w/v) uranyl acetate (Science Services, USA) in 70 % methanol for 10 min, followed by 0.4 % (w/v) lead citrate (Science Services, USA) in double-distilled water for 5 min. In addition, colloidal gold (20 nm diameter, BBI, UK) was attached to the semi-thick sections to serve as fiducial markers for the calculation of electron tomograms.

#### TEM of thin sections and pre-screening of semi-thick sections

To inspect thin sections, grids were analyzed by using a transmission electron microscope (Morgagni, Thermo Fisher) operated at 80 kV and equipped with a 2k x 2k CCD camera (Veleta, EMSIS). Thin sections were screened for developing and mature sperm cells. Semi-thick serial sections were also pre-screened using the same TEM to map regions of interest for subsequent 3D reconstruction by electron tomography.

#### Electron tomography and 3D reconstruction

Electron tomography was performed by using a transmission electron microscope (Tecnai F30, Thermo Fisher) operated at 300 kV and equipped with a 4k x 4k CMOS camera (OneView, Gatan). Using a dual-axis specimen holder (Type 2040, Fishione, USA), tilt series were recorded from -60° to +60° with 1° increments at a magnification of 4700 x and a pixel size of 2.572 nm applying the SerialEM software package(32,33). For dual-tilt electron tomography, the grid was rotated for 90° in the XY-plane and the second tilt series was acquired using identical microscope settings(34). Resulting tomographic A- and B-stacks of the same positions were reconstructed, combined and flattened using IMOD(35,36).

Serial-sections of the nebenkern were stitched and combined(37,38) by using automatically segmented microtubules as initial landmarks(39,40) within the ZIB Amira (Zuse Institute Berlin, Germany) software package(41). To automatically segment the membranes in the stitched serial tomograms of the nebenkern image processing, operations were performed within ZIB Amira that included the following computational operations: Morphological Laplacian (3D, 12 px, precision ‘faster’), Gaussian Filter (3D, separable, SD 5×5×5 px, kernel size factor ‘3’), Grayscale Fill Holes (3D, neighborhood ‘18’), Adaptive Histogram Equalization (3D, contrast limit ‘3’), and Adaptive Thresholding (3D, window 200×200×30 px, threshold ‘75’, criterion ‘greater-or-equal’, threshold mode ‘additive’). The binary image stack output was then used to generate a 3D surface. The same image processing pipeline was applied also to the single-section tomograms to segment membranes and chromatin. The central elements of the polycomplexes and the microtubules of the basal body were manually segmented. The microtubules of the axoneme and all other microtubules were automatically segmented using the ZIB Amira software package. Movies were also rendered using this software.

## Results

### Ultrastructure of spermatocytes in *H. membranacea*

The testes of *H. membranacea* were observed to have a large number of follicles, and individual testicular follicles could be observed macroscopically. The combination of ultrarapid high-pressure freezing after the dissection, freeze substitution with 1 % osmium tetroxide and 0.1 % uranyl acetate yielded an excellent preservation of the ultrastructure of the testes tissue for electron microscopy.

In each cell identified as being in prophase I we observed a nucleus surrounded by mitochondria. Golgi and ER could often be observed at the periphery of the cells (Fig. 1A-B). The nucleolus was often very prominently visible as dark stained structure inside of the nucleus (Fig. 1B-C). In prophase I, paired homologous chromosomes were observed with the central element of the synaptonemal complex (SC) visible as a thin line in between the aligned chromatin (Fig. 1C). We also observed multiple synaptonemal complexes stacked upon each other, structures called ‘polycomplexes’ (Fig. 1C; (42)). Electron tomography of a prophase nucleus showed that within one nucleus there exist regions with a single central element of the SCs and others with two or three (Fig. 1D, Movie 1). We measured the diameter of the SC to be 120 – 130 nm and found that the periodicity in between the stacked central elements within the polycomplexes was 110-120 nm. In further developed spermatocytes, we observed cells in prometaphase II in which the nuclear envelope has been broken down and chromosomes showing a high level of condensation (Fig. 1E). The mitochondria were arranged asymmetrically around the chromosomes in these cells. We also noticed ER or nuclear envelope remnants at the cell periphery and membrane-less protein aggregates of unknown origin or function (Fig. 1E, arrowheads). It appeared evident that the accumulation of mitochondria begins to be asymmetric within later meiotic stages (Fig. 1F). In some instances, we observed that the nuclear envelope is incomplete or not completely reformed (Fig 1F, arrowheads). Between the individual testicular follicles, we observed a connective tissue of multiple cell layers separating them from one another (Fig. 1G).

**Figure 1.**
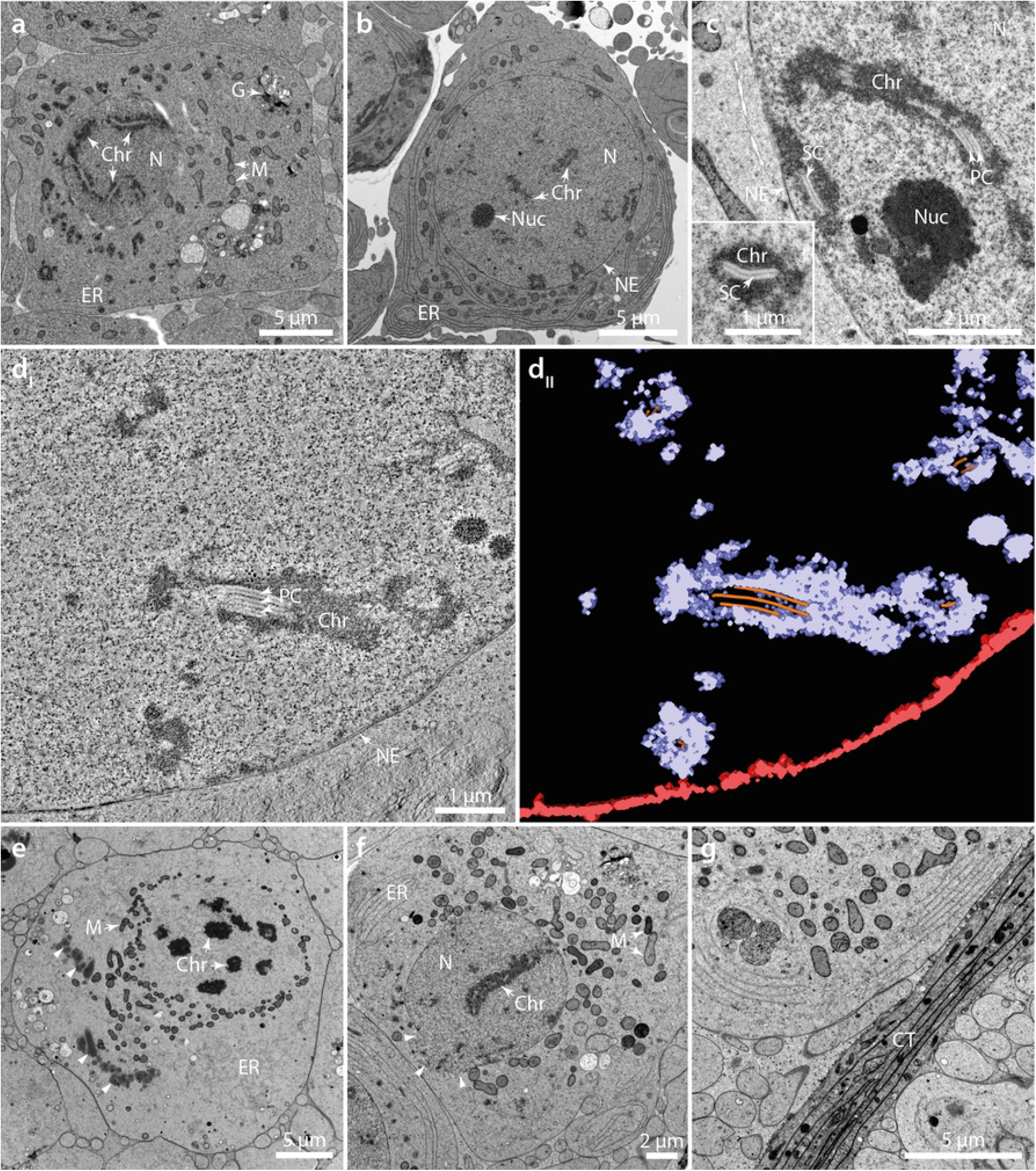
Ultrastructure of early spermatocyte development in *H. membranacea*. (**A**) TEM image of a spermatocyte in prophase of meiosis I. The nucleus (N) exhibits condensed chromatin (Chr) with synaptonemal complexes. Other organelles like the Golgi apparatus (G), the endoplasmic reticulum (ER) and mitochondria (M) are positioned in the vicinity of the nucleus. (**B**) Spermatocyte in prophase I. The nucleus (N) with a visible nuclear envelope (NE), a nucleolus (Nuc) and paired homologous chromosomes (Chr) are shown. The endoplasmic reticulum (ER) is positioned adjacent to the nucleus. (**C**) High-magnification view of a nucleus with condensed chromatin and a nucleolus. The central elements of synaptonemal complexes appear as single, double or triple complexes, a phenomenon known as polycomplexes (PC). The inset shows a synaptonemal complex at higher magnification. (**D**_**I**_) Tomographic slice of a three-dimensional reconstruction of a nucleus in prophase I. In this semi-thick section, the chromatin is positioned around multiple synaptonemal complexes and polycomplexes. (**D**_**II**_) Three-dimensional model of the nuclear envelope (red), the chromatin (purple) and central elements of the polycomplexes (orange). Same cell as shown in (D_I_). (**E**) TEM image of a cell at prometaphase I. The chromosomes are condensed and the nuclear envelope is broken down. Mitochondria surround the chromosomes. The ER and an electron-dense membrane-free aggregates (arrowheads) are visible in the cytoplasm. (**F**) TEM image of a spermatocyte presumably in meiosis II. The nuclear envelope appears open (arrow heads). (**G**) TEM of connective tissue (CT) separating the individual testicular follicles. The images are not shown according to the correct sequence of the developmental stages.

### Formation of the nebenkern

When investigating the TEM images of the testes section, we noted the existence of the nebenkern in spermatids, which is derived from fused mitochondria. This organelle later matures while wrapping around the axoneme, thereby providing the necessary energy for the movement of the sperm tail in mature sperm cells(20,43). Investigation of spermatids by phase contrast light microscopy revealed a dark, crescent shaped structure next to the nucleus (Fig. 2A). In later stages, this structure appeared more rounded and exhibited internal features (Fig. 2B), which resembles the structures we observed by TEM. In the later stages, it appeared less dense as observed with phase contrast light microscopy and internal membrane structures were more apparent (Fig. 2C). In some instances, we observed a dark structure at the nucleus that the pointed end of the nebenkern seemed to be pointing to. We suggest that this is a centriole adjunct (Fig. 2C-D), which tethers the basal body to the nucleus and provides structural support for the outgrowing axoneme. We also found a long dark structure spanning the distance from the nucleus to the cell membrane, which was surrounded by the very elongated nebenkern (Fig. 2D). We speculate that this structure is the axoneme, and the cell is about to grow the sperm’s tail.

**Figure 2.**
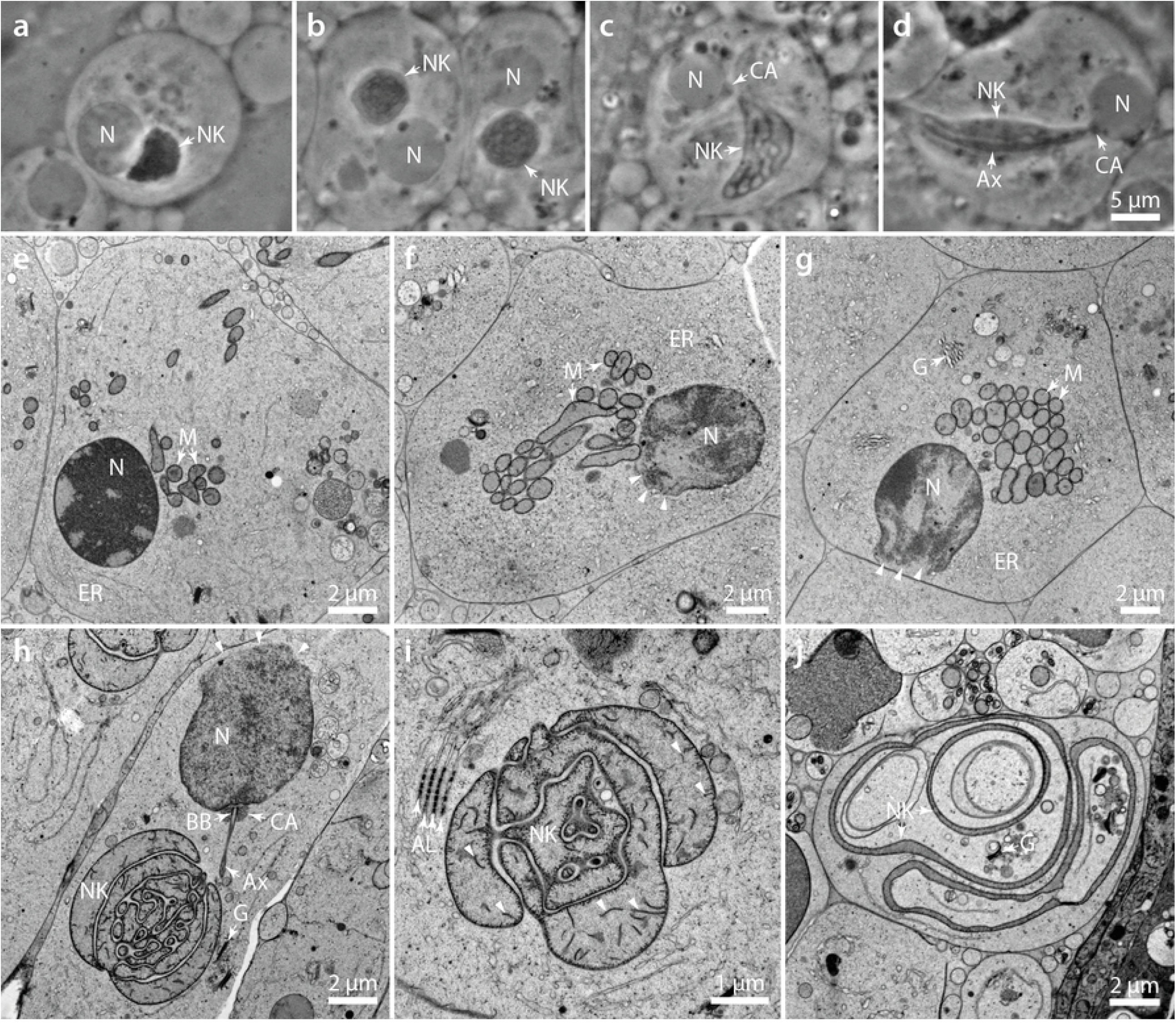
Fine structure of differentiating sperm cells. (**A**) Phase contrast image of a spermatid in an early stage of sperm maturation. The nucleus (N) and an irregular shaped nebenkern (NK) are shown. (**B**) Phase contrast image of a maturing spermatocyte. The nebenkern shows internal structures that cannot be resolved by phase contrast light microscopy. (**C**) Second example of a matured spermatocyte with an elongated, crescent-shaped nebenkern. Adjacent to the nucleus, a dark structure is visible that most likely represents a centriolar adjunct (CA) connecting the basal body to the nuclear envelope. (**D**) Spermatocyte with an axoneme (Ax) connected by the centriole adjunct material to the nucleus. The nebenkern appears elongated and spanning the whole volume of the cell. Scale bar for (A-D), 5 µm. (**E**) TEM image of a cell in late telophase II. The chromatin is still densely packed inside the nucleus, which is surrounded by endoplasmic reticulum (ER). Mitochondria (M) start to aggregate at a position close to the nucleus. (**F-G**) Sperm cell with clustered mitochondria, ER and Golgi (G). The nucleus (N) shows a fenestrated nuclear envelope (arrow heads). This fenestration is likely to be involved in a shrinkage and compaction of the nucleoplasm. (**H**) Nebenkern and nucleus with an attached basal body (BB) anchored through the centriolar adjunct structure (CA). The developing axoneme and the Golgi apparatus are visible in the vicinity of the nebenkern. The nuclear envelope appears fenestrated (arrowheads) at the opposite side of the nebenkern. (**I**) Maturing of the nebenkern into a complex multi-membrane structure. The cristae of the former mitochondria are still visible within the lumen of the nebenkern (arrowheads). Annulate lamellae (AL) are also visible presumably on an ER-derived structure next to it. (**J**) Nebenkern after maturation. In later sperm development, the nebenkern unfolds and assembles in the cytoplasm around the axoneme. In between this structure there are still cytoplasm and organelles like Golgi (G) visible.

By TEM we observed that in telophase the mitochondria begin to cluster together actively at one side of the nucleus (Fig. 2E). This clustering becomes very prominent with progressing cell maturation (Fig. 2F-G). We believe that the mitochondria start to fuse together at the time of dense clustering. Interestingly, in these cells we always observed an open nuclear envelope. Presumably, the nucleus shrinks in volume at this time and that this opening then serves to release the nucleoplasm (Fig. 2F-G, arrow heads). In further developed cells, the mitochondria have fused largely to form the nebenkern, a complex three-dimensional (3D) arrangement of membranes and is always close to the nucleus to which the basal body is anchored. This attachment is achieved by a granular, amorphous structure described as the centriole adjunct (Fig. 2H).

From the basal body we observed the outgrowing axoneme in the direction of the nebenkern. In some instances, we also saw annulate lamellae in the male germ cells, an ordered structure of nuclear pore complexes within the endoplasmic reticulum and hence close to the nucleus or nebenkern (Fig. 2I). Although the luminal volume of the nebenkern increases with more fusion events happening, there are still cristae visible inside within in the lumen itself as well as adjacent to the membrane (Fig. 2I, arrow heads). In further developing cells the nebenkern changed its shape and became a multilayered, very elongated membrane organelle, whose shape can be hardly described by analyzing single sections (Fig. 2J). We noticed that in between the multiple membrane layers there are cellular organelles like Golgi and, therefore, the structure seems to fold around through the cytoplasm. In our understanding, the TEM images shown in Fig. 2 H-I correspond to the light microscopic image shown in Fig. 2B, and the TEM image in Fig. 2J corresponds to Figs. 2 C-D.

### Three-dimensional reconstruction of the matured nebenkern

Previous work showed that the nebenkern consists of two independent but interlocked entities^20^ as observed in the harlequin cabbage bug *Murgantia histrionica*(17). Therefore, we sought to apply electron tomography to analyze the three-dimensional (3D) architecture of the nebenkern in *H. membranacea*. We acquired electron tomograms from seven serial semi-thick (300 nm) sections (Fig. 3A_I-IV_). By stitching these serial tomograms, we were able to cover about one fourth of the volume of a nebenkern (Fig. 3B_I-IV_). After segmentation of the membranes in the tomographic sections, we then rendered the surfaces to obtain a 3D model of the nebenkern (Fig. 3C, Movie 2). We then displayed the inner volume of the nebenkern by clipping it along either the x-(Fig. 3D, Movie 3) or the y-axis (Fig. 3E, Movie 4) or both (Fig. 3F). We observed that within the lumen of the nebenkern there were numerous membrane invaginations resembling the cristae of mitochondria. By further analyzing the 3D model we observed that the cytoplasm that appeared trapped within the nebenkern volume at a first glance was in fact not separated from the cytoplasm of the whole cell as it was not entirely enclosed by membranes.

**Figure 3.**
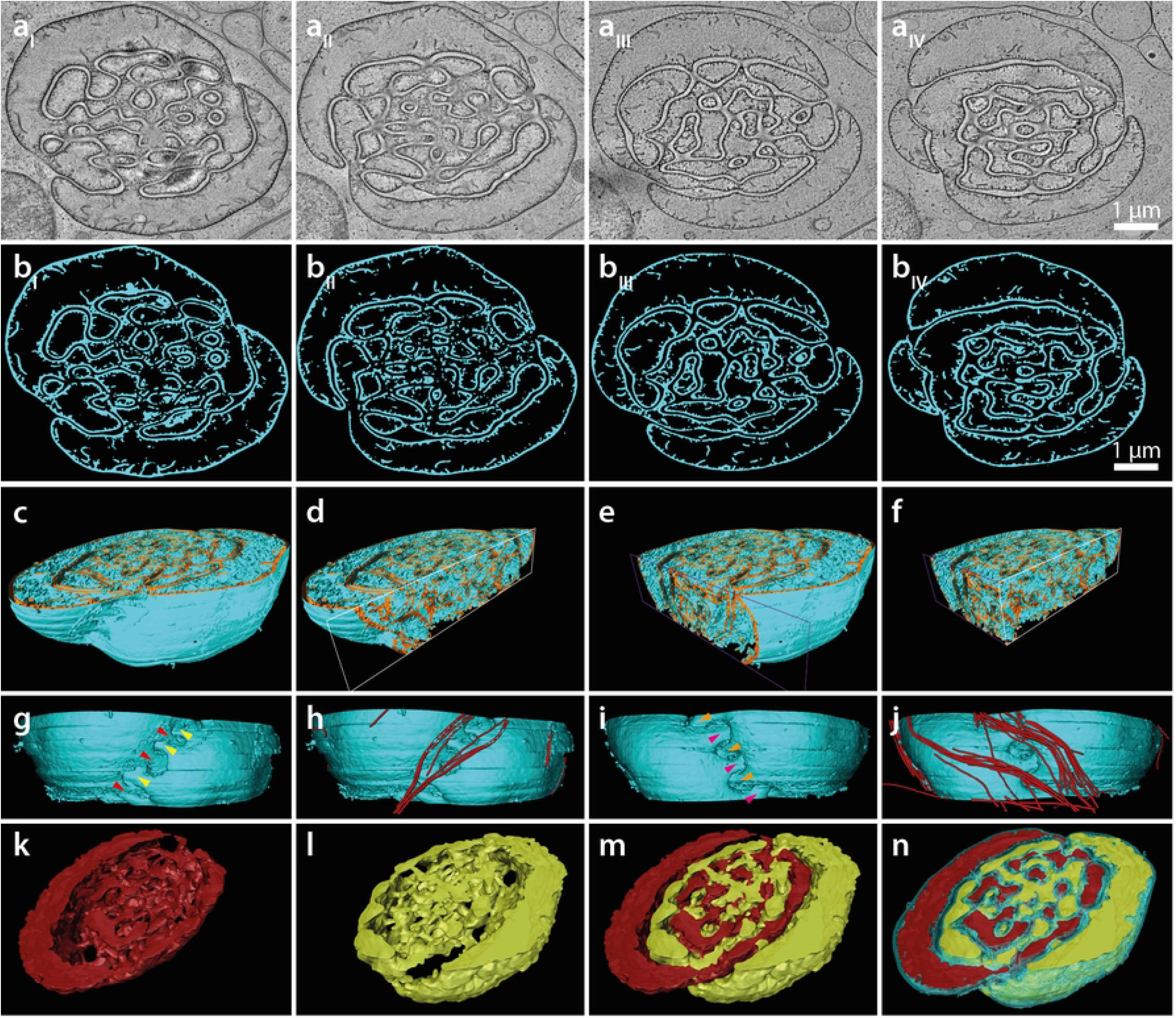
Partial 3D reconstruction of the nebenkern covering about 2.1 µm in the z-dimension. (**A**_**I**_**-A**_**IV**_) Tomographic slices through a nebenkern. (**B**_**I**_**-B**_**IV**_) Segmentation of membranes (cyan) in the tomographic slices as shown in (A). (**C-F**) Side views of the 3D model of the nebenkern with two clipping planes (x-axis white, y-axis purple) perpendicular to each other, illustrating the interlocking of the internal membranes of the nebenkern. The membrane is shown in cyan and then membrane lumen of the nebenkern is shown in orange. (**G**) Front view of the 3D model of the nebenkern illustrating a zipper-like membrane structure. The arrow heads (differently colored) illustrate the direction of the membrane extrusions leading into the nebenkern from opposite positions. (**H**) Identical view showing microtubules (red) in close proximity of the membrane structure. (**I-J**) Back view on the nebenkern model with respective microtubule model. (**K**) Segmentation of the interconnected inner lumen of one out of two mitochondria-derived nebenkern constituents (shown in red). (**L**) Segmentation of the other mitochondria-derived nebenkern constituents (shown in yellow). (**M**) 3D model of both nebenkern constituents illustrating the interlocking of both membrane systems. (**N**) 3D model of the nebenkern with the membrane (cyan) and both segmented inner lumina of the constituents (red and yellow).

In addition, we also observed two zipper-like membrane structures positioned directly opposite one another in our nebenkern reconstruction (Fig. 3G-J, Movie 5). This zipper-like structure was not apparent when looking at single thin sections, emphasizing again the importance of 3D imaging for a spatial analysis of organelle organization. The membranes seemed to be connected in an alternating fashion (Fig. 3G, I, arrow heads). We also observed an accumulation of microtubules at this region indicating a possible involvement of the cytoskeleton in this process (Fig. 3H, J, Movie 5).

Most importantly, however, our analysis revealed that there are two membrane entities that are inter-connected but both membrane systems never touched each other in the volume that has been investigated (Fig. 3K-N, Movie 6). The ‘zipper-like’ structure at the outside is the region where both halves of the nebenkern are interwoven.

### Sperm maturation

In testicular follicles containing more mature sperm we observed axonemes next to slightly more electron-dense, membrane structures showing membrane extrusions within their lumen which is presumably the matured nebenkern (Fig. 4A-B). These structures were often obvious as round objects connected by a thin tube-like connection. In some instances, we found multiple axonemes next to each other (Fig. 4A, arrow heads). In the investigated regions we also observed Golgi and thin membrane systems, which might be ER or its remnants. In some instances, we found that the nuclear envelope that is in contact with the centriole adjunct is densely stained thus appearing thickened (Fig. 4C, arrowheads). The anchor that tethers the basal body to the nuclear envelope consisted of small darkly stained dots that resemble ribosomes in TEM images (Fig. 4C). This motivated us to investigate this structure in detail using electron tomography. Within the reconstructed tomogram we saw that the nuclear envelope was in fact not thickened but appeared clearly bi-layered, with a darker staining at the outside, facing the centriole adjunct (Movie 7). These darkly stained structures within the centriole adjunct were 15-20 nm in diameter and appeared to be interconnected in a “beads-on-a-string” fashion (Movie 7).

**Figure 4.**
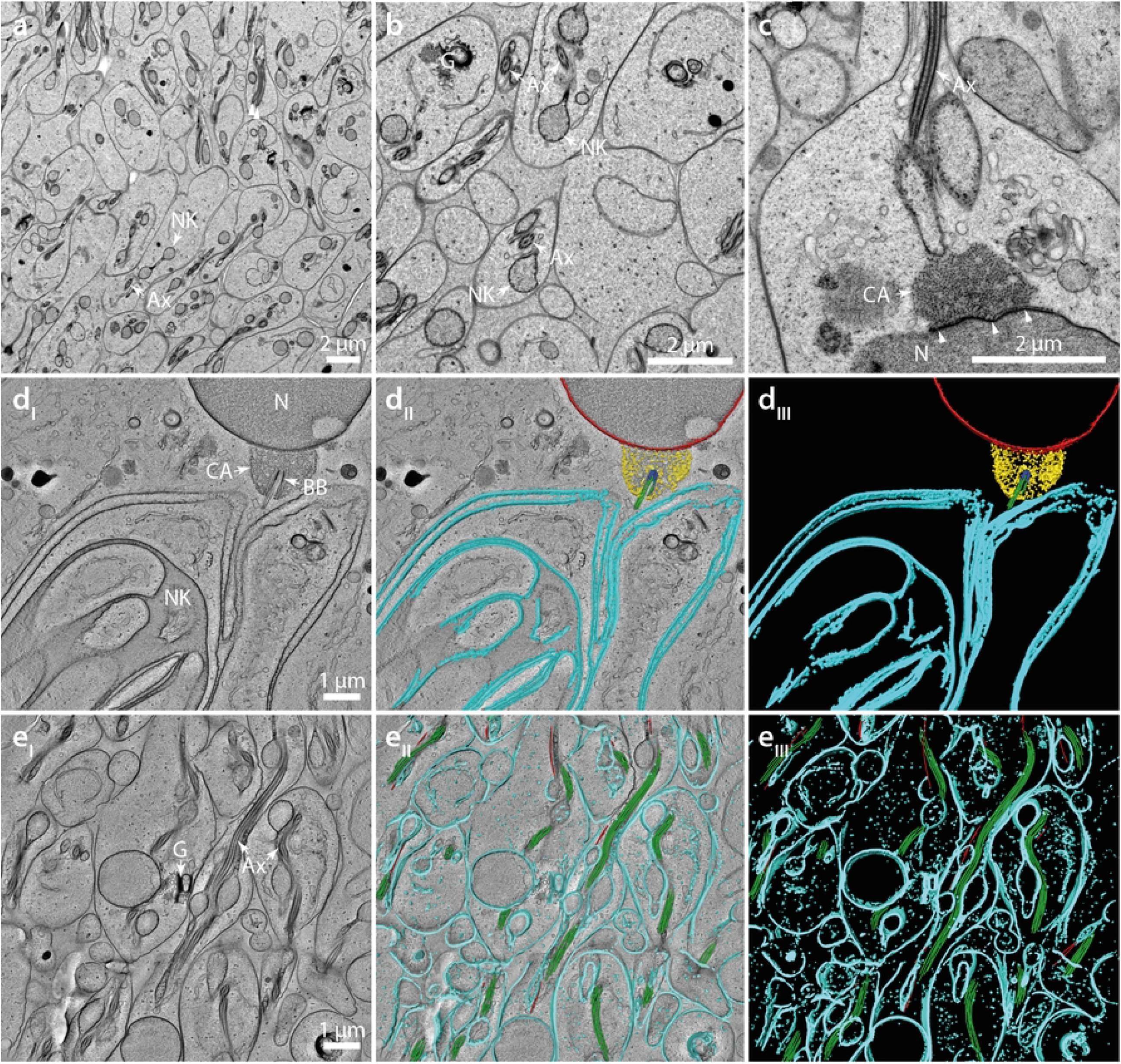
Ultrastructure of spermiogenesis in *H. membranacea*. (**A**) Thin-section-TEM image of a testes region. Multiple developing sperm cells are visible with axonemes (Ax) and nebenkern derivatives (NK). (**B**) Detailed view of a region with multiple Golgi apparati (G), axonemes (Ax) and nebenkern derivatives (NK) visible. (**C**) High-magnification view of a sperm cell illustrating the granular character of the centriolar adjunct (CA) next to the nucleus (N). An electron-dense structure is indicated (arrow heads). An outgrowing axoneme (Ax) is visible at an opposing position. (**D**_**I**_**-D**_**III**_) Electron tomogram and segmentation of the nucleus with an attached basal body (BB). The centriolar adjunct surrounds the basal body with the outgrowing axonemal microtubules. A developing nebenkern is visible next to the axoneme. The nuclear envelope is shown in red, the nebenkern membrane in cyan, the centriole adjunct in yellow, the basal body in blue and outgrowing microtubules in green. (**E**_**I**_**-E**_**III**_) Electron tomogram and segmentation of a region containing developing sperm tails. The Golgi apparatus and many axonemes are shown in cross- and longitudinal sections. In the 3D models, membranes are illustrated in cyan, axonemal microtubules in green and other microtubules in red.

Tomographic reconstruction also revealed that the axoneme grows in the direction of the nebenkern which forms a pouch presumably resembling an opening or the bipartite split of the two nebenkern-derivatives that engulfs the axoneme (Fig. 4D_I-III_, Movie 8). In an area with more mature sperm we acquired another tomogram in order to illustrate that the nebenkern next to the axoneme consists indeed of round structures that are connected by a tube-like membrane structure (Fig. 4E, Movie 9). They were always in close proximity to the axoneme and exhibited another set of microtubules in their cytoplasm that were clearly different from axonemal microtubules, as well as Golgi stacks (Fig. 4E, Movie 10).

## Discussion

By applying high-pressure freezing in combination with 3D reconstruction by serial-section electron tomography, we report on the ultrastructure of the nebenkern and other cytological features during spermatogenesis in the praying mantid, *Hierodula membranacea*. In previous studies, the ultrastructure of male germ cells of other insect species was investigated in chemically fixed samples(44). This classical method of fixation suffers from a slow rate of diffusion of the fixative into cells and tissues(44). Depending on the sample and the occurrence of cuticles and other mechanical barriers, the diffusion rate of the fixative is less than one mm per hour and therefore can take up to minutes or several hours(45). To avoid any preparation artifacts, we performed ultrarapid cryo-immobilization followed by freeze substitution. Applying this approach, we achieved an excellent ultrastructural preservation.

Visualizing spermatids in a range of stages of maturation, the ultrastructure of spermatogenesis in *H. membranacea* at the first glance is similar to that observed in other insects(12). Unexpectedly, however, we observed multiple synaptonemal complexes in stacks in areas within a single chromosome (Fig. 1C), a phenomenon previously described as ‘polycomplexes’(42). We noted the existence of polycomplexes in prophase I. This structure has been documented as well in other eukaryotes (e.g. *D. melanogaster, S. cerevisia*e and *C. elegans*)(42).

An additional interesting observation is the plasticity and complexity of the nebenkern formation. Until the end of meiosis II, multiple individual mitochondria are visible in the vicinity of the nucleus. In telophase II, there is an apparent accumulation of the mitochondria close to the nucleus. Although we do not have data on the dynamics of this process, we speculate that this mitochondrial accumulation is achieved through an association with microtubule. Support for this hypothesis might be provided by a detailed spatial analysis of microtubules in the vicinity of mitochondria. By addition of nocodazole to isolated spermatocytes, one could also perturb the cytoskeleton to test whether a disruption of microtubule dynamics might prevent the formation of the nebenkern.

A puzzling aspect of nebenkern formation is how the mitochondria fuse to form such an enormous organelle. Mitochondrial fusion is known to be mediated by the GTPases mitofusin-1 and mitofusin-2 (*Mfn1, Mfn2*) which are located in the outer mitochondrial membrane(23,46). In *D. melanogaster* it has been shown that fuzzy onions (fzo) mutants show defects in nebenkern formation and lead to sterility(22). The question arises, do processes that are involved in the nebenkern formation in *H. membranacea* also rely on this mitochondrial fusion machinery? To test this, either RNAi and/or CRISPR/Cas technology would need to be developed in this species to inhibit mitofusin activity. In addition, the maturation of the nebenkern is a fascinating cell biological topic. After a labyrinth-like state of the nebenkern is formed (Fig. 2H-I, Fig. 3), this organelle completely flattens out to adopt a layered structure composed of thin sheets (Fig. 2J), culminating in a tube formation (Fig. 4A-B, E). The 3D organization and the molecular mechanisms of this morphological process also remain to be unraveled.

By segmenting the inner lumen of the nebenkern and grouping of connected parts we showed that the nebenkern at this stage is indeed composed of two parts (Fig. 3K-N). The complex nature of the interlocked segments and the fact that both parts did not seem to touch each other raises the question of how the two parts are built in the first place and how the partitioning then is achieved. Further structural studies of earlier and later nebenkern stages are clearly needed to better understand the morphological changes that take place during partitioning and subsequent flattening of this organelle.

Interestingly, we detected a zipper-like membrane arrangement on two opposing sides of the nebenkern (Fig. 3G-J, Movie 5). This membrane shape was not obvious when looking at the segmented, stitched z-planes (Fig. 3B). Only rendering of the membrane made it possible to analyze the 3D organization. In *Drosophila melanogaster* and *Murgantia histrionica*, the data suggest that the nebenkern splits into two derivatives, finally wrapping around the axoneme in the sperm tail(17,20,43). If this holds true also for *H. membranacea*, it would explain the observed two-fold symmetry of this structure and the observed ‘zipper-like’ structure could be a result of this process. The presence of microtubules on both ‘zipper-like’ membrane structures could indicate the involvement of the microtubule cytoskeleton and motor proteins in the separation or movement of this structure in the cell.

Our 3D electron microscopic analysis revealed that the nebenkern in *Hierodula membranacea* is indeed composed of two interwoven segments that are connected by a ‘zipper-like’ structure at opposing positions. Our ultrastructural approach could be applied to an analysis of spermatogenesis also in other insect species with similar structures to investigate if these features of nebenkern formation are evolutionarily conserved.

## Acknowledgements

We would like to thank the members of the Core Facility Cellular Imaging (CFCI, Faculty of Medicine Carl Gustav Carus, TU Dresden) for help with light microscopy and the EM facility at MPI-CBG, Dresden for technical assistance with electron tomography. We would also thank various members of the slack channel “EMofCellsTissuesOrganisms” for help with identifying structures. We are also grateful to Dr. Daniel Baum (Zuse Institut Berlin) for support in using ZIB Amira. This work was supported by grants from the National Science Foundation (NSF, RUI-1715157 to L.P.) and the Deutsche Forschungsgemeinschaft (DFG, MU1423/10-1 to T.M-R.).

## Movie legends

**Movie 1**. Electron tomographic reconstruction of a single semi-tick (300 nm) section of a nucleus in prophase I. The nuclear envelope (red), chromatin (purple), and the central elements (orange) of synaptonemal complexes and polycomplexes are shown. This movie corresponds to Fig. 1D.

**Movie 2**. Movie of seven registered and stitched electron tomographic reconstructions of the nebenkern covering about 2.1 µm in the z-dimension. The membrane of the nebenkern is shown in cyan, the membrane lumen in orange. This movie corresponds to Fig. 3A-C.

**Movie 3**. Stitched and segmented nebenkern with a consecutive clipping along the x-axis. This 3D movie illustrates the structural complexity of the nebenkern. The membrane of the nebenkern is shown in cyan, the membrane lumen in orange. The clipping plane is indicated by a white rectangle. This movie corresponds to Fig. 3D.

**Movie 4. S**titched and segmented nebenkern with a consecutive clipping along the y-axis. This 3D movie illustrates the structural complexity of the nebenkern. The membrane of the nebenkern is shown in cyan, the membrane lumen in orange. The clipping plane is indicated by a purple rectangle. This movie corresponds to Fig. 3E.

**Movie 5**. Side view of the stitched and segmented nebenkern, illustrating a zipper-like membrane structure at both sides of the nebenkern. The membrane of the nebenkern is shown in cyan. Microtubules (red) can be observed adjacent to the nebenkern membrane. This movie corresponds to Fig. 3G-J.

**Movie 6**. Segmentation of the interconnected inner lumen of the two mitochondria-derived nebenkern constituents (shown in red and yellow). The membrane is shown in cyan. This movie corresponds to Fig. 3K-N.

**Movie 7**. Electron tomographic reconstruction of a single semi-tick (300 nm) section of a nucleus with an attached basal body and the centriole adjunct. This movie corresponds to Fig. 4D and Movie 8.

**Movie 8**. Electron tomographic reconstruction of a single semi-tick (300 nm) section of a nucleus with an attached basal body. The centriole adjunct (yellow) surrounds the basal body (blue) with outgrowing axonemal microtubules (green). The nuclear envelope is shown in red, the developing nebenkern in cyan. This movie corresponds to Fig. 4D and Movie 7.

**Movie 9**. Electron tomographic reconstruction of a single semi-tick (300 nm) section of a region containing the developing axonemes and multiple nebenkern structures. Axonemes in various orientations can be seen. Membranes are shown in cyan, axonemal microtubules in green and other microtubules in red. This movie corresponds to Fig. 4E and Movie 10.

**Movie 10**. Detailed view of an electron tomographic reconstruction of a single semi-tick (300 nm) section of a region showing the developing axonemes and multiple nebenkern structures. This movie corresponds to Fig. 4E and Movie 9.

